# Tissue volume estimation and age prediction using rapid structural brain scans

**DOI:** 10.1101/2022.01.19.476615

**Authors:** Harriet Hobday, James H. Cole, Ryan A. Stanyard, Richard E. Daws, Vincent Giampietro, Owen O’Daly, Robert Leech, František Váša

## Abstract

The multicontrast EPImix sequence generates 6 contrasts, including a T_1_-weighted scan, in ∼1 minute. EPImix shows comparable diagnostic performance to conventional scans under qualitative clinical evaluation, and similarities in simple quantitative measures including contrast intensity. However, EPImix scans have not yet been compared to standard MRI scans using established quantitative measures. In this study, we compared conventional and EPImix-derived T_1_-weighted scans of 64 healthy participants using tissue volume estimates and predicted brain-age. All scans were pre-processed using the SPM12 *DARTEL* pipeline, generating measures of grey matter, white matter and cerebrospinal fluid volume. Brain-age was predicted using *brainageR*, a Gaussian process regression model previously trained on a large sample of standard T_1_-weighted scans. Estimates of both global and voxel-wise tissue volume showed significantly similar results between standard and EPImix-derived T_1_-weighted scans. Brain-age estimates from both sequences were significantly correlated, although EPImix T_1_-weighted scans showed a systematic offset in predictions of chronological age. Supplementary analyses suggest that this is likely caused by the reduced field of view of EPImix scans, and the use of a brain-age model trained using conventional T_1_-weighted scans. However, this systematic error can be corrected using additional regression of T_1_-predicted brain-age onto EPImix-predicted brain-age. Finally, retest EPImix scans acquired for 10 participants demonstrated high test-retest reliability in all evaluated quantitative measurements. Quantitative analysis of EPImix scans holds potential to reduce scanning time, increasing participant comfort and reducing cost, as well as to support automation of scanning, utilising active learning for faster and individually-tailored (neuro)imaging.

## Introduction

### EPImix

A new multicontrast magnetic resonance (MR) pulse sequence has been developed, named EPImix (Skare et al., 2018). This sequence utilises single-shot echo-planar imaging (EPI) and different magnetisation preparations to rapidly generate 6 contrasts; T_1_-FLAIR, T_2_-FLAIR, T_2_-weighted, T_2_*-weighted, diffusion-weighted imaging (DWI) and apparent diffusion coefficient (ADC) (for full acquisition details, see Skare et al. (2018)). The EPImix sequence has lower signal-to-noise ratio and resolution than standard MRI sequences (Skare et al., 2018), resulting in lower image quality ratings than routinely collected corresponding MR scans (Delgado et al., 2019; Ryu et al., 2020). However, the main benefit of this single-shot, multicontrast sequence over routinely acquired individual MRI sequences is its speed. The EPImix sequence is faster than corresponding single-contrast sequences because of its lower matrix size and because an EPI readout is more SNR efficient than analogous FSE readouts (Skare et al., 2018). This multimodal sequence can acquire full brain coverage in 78 seconds, relative to the ∼750 seconds needed to collect these 6 contrasts using standard MRI sequences (Delgado et al., 2019; Ryu et al., 2020). In addition to being more cost effective (Mekle et al., 2006), shorter scanning times improve participants’ comfort and reduce motion (Andre et al., 2015), potentially resulting in less unusable data, especially for clinical populations (Greene et al., 2016). The EPImix sequence also generates multiple types of MR contrasts, therefore capturing more tissue characteristics than standard parametric mapping (Delgado et al., 2019; Ryu et al., 2020).

### Quantitative analyses

To establish whether this new sequence holds potential to reduce scanning time whilst producing similar derived measures, it needs to be quantitatively^1^ compared to corresponding routinely acquired MR scans. Previous research established that EPImix scans are comparable to conventional scans in the context of qualitative clinical diagnosis (Delgado et al., 2019; Ryu et al., 2020). Moreover, our previous work compared EPImix and routine T_1_-weighted (T_1_-w) scans^2^ using simple rapidly-derived quantitative measurements, including image intensity and Jacobian determinants (obtained from the registration to MNI standard space), and found high similarity in these rudimentary quantitative measures as well as their potential to identify inter-individual differences (Váša et al., 2021). We further demonstrated the utility of the multicontrast EPImix sequence to construct morphometric similarity networks (MSNs), which provide individual estimates of anatomical connectivity (Seidlitz et al., 2018), in minutes following participants entering the MRI scanner (Váša et al., 2021). However, EPImix scans have not previously been compared to standard sequences using established quantitative structural neuroimaging measures such as estimates of tissue volume (Lerch et al., 2017) or predicted brain-age, a putative biomarker of brain health (Cole and Franke, 2017; Cole et al., 2019b). Such quantitative measures likely reflect subtle inter-individual differences and provide a more thorough comparison between EPImix and standard sequences than qualitative measures (Pierpaoli, 2010) or potentially noisy estimates such as tissue intensity (Fortin et al., 2016). Moreover, the ability to derive commonly used quantitative measures from data acquired with faster imaging sequences, such as EPImix, has the potential to reduce scanning time, increasing participant comfort and reducing costs.

### Active Acquisition

One potential application of the EPImix sequence is in active acquisition. Active acquisition is a proposed type of data acquisition which would utilise active learning (Settles, 2009), to analyse MRI data as it is acquired, with results used to guide further image acquisition, in a closed-circuit sequence (Lorenz et al., 2016; Cole et al., 2019a). This would remove the necessity of making a priori decisions about scanning parameters – such as the type of scan, scan resolution and/or the scan location. Instead, these would be dependent on the individual inside the scanner, driving the selection of scanning sequences towards identification of individual differences or personalised clinical diagnosis. In this way, active acquisition holds the potential for reduced timing and improved accuracy, reliability and individualisation of (neuro)imaging (Cole et al., 2019a). The main feasibility obstacles to active acquisition are the speed of image collection as well as data processing and analysis; to realise the benefits of this multimodal adaptive approach, analyses need to be carried out in near real-time (Cole et al., 2019a; Váša et al., 2021). Due to the relative speed of the new EPImix sequence, it could contribute considerably to this process.

### Quantitative comparison of EPImix and standard T_1_ contrasts

Here, we focused on the T_1_-w EPImix contrast and compared it to a standard T_1_-w contrast at two levels. We first compared T_1_-w global and voxel-wise estimates of grey matter (GM), white matter (WM) and cerebrospinal fluid (CSF) tissue volumes (Lerch et al., 2017). These tissue volumes vary as a function of age and are typically abnormal in many psychiatric and neurological populations (Ramanoël et al., 2018; Wang et al., 2019).

We then compared predicted brain-ages from EPImix and standard T_1_-w scans to assess whether EPImix can reliably detect the well-established relationship between healthy aging and changes in tissue volumes (Hafkemeijer et al., 2014; Ramanoël et al., 2018). Predicted brain-ages are derived from a validated multivariate regression model that estimates the biological age of an individual brain directly from the imaging data. This estimate provides a useful summary of the large voxelwise spaces generated by standard high-resolution T_1_-w scans into a single interpretable value. The difference between the predicted brain-age and the true chronological age is a versatile measure that has been associated with cognitive ability (Cole et al., 2019b; Cole, 2020) as well as clinical status and severity, with significantly “older” brain-ages associated with traumatic brain injury (TBI) (Cole et al., 2015), mild cognitive impairment (MCI) and Alzheimer’s disease (AD) (Palma et al., 2020). Furthermore, predicted brain-age could be a better predictor of disease risk than chronological age (Franke and Gaser, 2019; Cole, 2020). For example, brain-age has been shown to be a reliable predictor of which individuals will develop AD (Gaser et al., 2013; Biondo et al., 2021).

We expected quantitative measurements derived from EPImix T_1_-w scans, including global and voxelwise estimates of tissue volume as well as predicted brain-age, to be broadly comparable to analogous estimates derived from standard T_1_-w scans. Given the notable differences in the field of view (FoV) between EPImix and standard T_1_-w scans, the tissue volume and brain-age analyses were repeated with standard T_1_-w scans that had their FoV artificially reduced to match the FoV of the EPImix T_1_-w contrast. Finally, we assessed the within-session test-retest reliability of the EPImix-derived estimates of tissue volume and predicted brain-age.

## Methods

### Participants

EPImix scans (including a T_1_-w contrast) were acquired for 95 healthy participants across three studies that used the same MRI scanner; one participant with particularly reduced cortical coverage (due to reduced FoV of EPImix scans, discussed below) was excluded, resulting in EPImix scans from 94 participants (F: 47, M: 47) used for analyses. The age range was 18-59 years. The mean age of participants was 28.1 years, with a standard deviation of 9.2 years (female mean age: 26.7 ± 7.1 years; male mean age 29.6 ± 10.7 years). Of those 94 participants, 64 (F: 32, M: 32) were also scanned using a standard high-resolution T_1_-w sequence. The age range of this subset of participants was 18-59 years; their mean age was 28.9 years, with a standard deviation of 10.1 years (female mean age: 22.6 ± 1.8 years; male mean age: 35.3 ± 11.2 years); for details, see Supplementary Information (SI) Fig. S1. All three studies received ethical approval from King’s College London’s Psychiatry, Nursing and Midwifery Research Ethics Committee (KCL Ethics References: HR-18/19-9268, HR-18/19-11058 and HR-19/20-14585). All participants gave written informed consent to take part in their respective study.

### MRI acquisition

All scans were collected on the same General Electric (GE) MR750 3T scanner (Waukesha, WI). EPImix scans were acquired from 94 participants, consisting of six contrasts (T_2_*, T_2_-FLAIR, T_2_, T_1_-FLAIR, DWI, ADC). Only the T_1_-FLAIR contrast was used in this study (referred to as the EPImix T_1_-w contrast), which was acquired with the following parameters: TE = 16.5 ms, TR = 1300 ms, TI = 582 ms, flip angle = 90°, matrix size = 180 × 180, FoV 240 mm, 32 slices, slice thickness = 3 mm, voxel resolution = 0.975 × 0.975 × 3 mm. The EPImix sequence includes an in-scanner motion correction step (Skare et al., 2018) and we used the motion-corrected images for analyses. For details regarding the EPImix sequence, see Skare et al. (2018). Additionally, for 10 participants a second EPImix scan was acquired during the same session, which was used to quantify test-retest reliability.

Additionally, conventional T_1_-w scans were acquired for 64 participants within the same session, using an IR-FSPGR sequence. Of these, 12 scans were acquired with the following parameters: TE = 3.172 ms, TR = 8.148 s, flip angle = 12°, matrix size = 256 × 256, FoV = 256mm, 164 slices, slice thickness = 1 mm, voxel resolution 1 × 1 × 1 mm; and 52 scans were acquired with the following parameters : TE = 3.016 ms, TR = 7.312 ms, flip angle = 11°, matrix size = 256 × 256, FoV = 270mm, 196 slices, voxel resolution = 1.05 × 1.05 × 1.2 mm.

### Pre-processing and brain-age estimation

All brain scans were pre-processed using a SPM12 *DAR-TEL* processing pipeline (Ashburner et al., 2014); including bias field correction, segmentation, registration to standard space (MNI152, 6th generation) via an intermediate study-specific template, and smoothing using a 4mm FWHM kernel. This process generated voxelwise estimates of tissue volumes, including grey matter (GM), white matter (WM), and cerebrospinal fluid (CSF). Global tissue volumes were calculated by summing across voxels in each tissue class.

The above processing pipeline was applied as part of the *brainageR* software, which was additionally used to obtain brain age predictions. The brain-age model was previously trained to predict chronological age from conventional T_1_-w scans in 3377 healthy people aged 18-92 years, using Gaussian Processes Regression. For additional information about the brain-age model, see Cole, 2019 and https://github.com/james-cole/brainageR.

The reduced FoV of the EPImix acquisition used in this study resulted in imperfect cortical coverage in some participants; in particular, portions of the inferior temporal and/or superior parietal lobes were not consistently scanned (SI Fig. S2). To investigate the potential impact of this reduced FoV on quantitative analyses, we repeated these using standard T_1_-w scans with artificially reduced FoV, to match the reduced EPImix FoV (within-participants). This was performed using affine registration (FSL FLIRT) (Jenkinson et al., 2002) of the standard T_1_-w scan to an upsampled version of the same participant’s EPImix T_1_-w contrast (resampled from the EPImix resolution of 0.975 ×0.975 × 3 mm, to match the standard T_1_-w resolution of 1 × 1 × 1 mm, or 1.05 × 1.05 × 1.2 mm). Standard T_1_-w scans with artificially reduced FoV (T_1_-w FoV_*EP I*_) were then processed using the steps described above, for both tissue volume estimation and brain age prediction.

Processing was run on a Dell workstation (16-CPU 3.6GHz Intel Xeon, 128Gb RAM). Total processing time was recorded for each scan, including data processing using SPM12 and brain-age estimation. The processing time of EPImix T_1_-w scans was: median [1st, 3rd Quartile] (Md [Q_1_,Q_3_]) = 4.82 [4.75, 4.9] min; for standard T_1_-w scans, Md [Q_1_,Q_3_] = 7.61 [7.50, 8.17] min; for standard T_1_-w scans with reduced FoV, Md [Q_1_,Q_3_] = 5.42 [5.37, 5.66] min (SI Fig. S3).

### Statistical analysis

Quantitative measures derived from EPImix T_1_-w scans, standard T_1_-w scans and T_1_-w scans with reduced FoV from 64 participants were compared using the Spearman’s correlation coefficient (*r*_*s*_). We first compared tissue volumes (for GM, WM and CSF) between sequences, both at the global and voxelwise levels. Voxelwise comparisons were restricted to voxels contained within the FoV of EPImix scans in the majority of participants (i.e. voxels with at least 0.001 mm^3^ tissue volume in at least 95% of participants). Furthermore, we quantified the relationship between GM volume and participant chronological age, to ascertain whether EPImix T_1_-w scans are equally sensitive as standard T_1_-w acquisitions to known decreases in GM volume in ageing (Hafkemeijer et al., 2014; Ramanoël et al., 2018).

We next compared estimates of predicted brain-age to true (chronological) age, across sequences, using Spearman’s correlation coefficient (*r*_*s*_), the proportion of variance explained (*r*^2^) as well as the median absolute error (MAE). We also compared the predicted brain-ages between scan types.

As the correspondence between brain age predictions from standard and EPImix T_1_-w scans was imperfect (likely due to the brain-age model being trained on standard T_1_-w scans; see *Results* & *Discussion*), we explored the use of an additional regression step to optimise brain age prediction from EPImix T_1_-w scans. We used leave-one-out cross-validation to regress T_1_-w predicted brain-age on EPImix-predicted brain-age (in 63 participants), leading to an adjusted estimate of predicted brain-age in the remaining (left-out) participant. Repeating this approach by iterating across all participants enabled us to quantify the MAE (relative to chronological age) of this adjusted brain-age estimate.

Finally, to benchmark the relative ability of each scan type (T_1_-w, EPImix T_1_-w, T_1_-w FoV_*EPI*_) to predict brainage, we compared each MAE estimate to the MAE from the worst possible brain-age model. Our “null model” MAE estimate was based on an assumed prediction of the same brain-age for every participant, equivalent to the mean age of participants within the training dataset of the *brainageR* model (40.6 years); this led to a null MAE of 15.6 years.

We then calculated the ratio of this null MAE to the MAE of each scan type, to estimate the extent of predictive improvement of each scan type relative to the worst possible model.

Most analyses were carried out using the sample of 64 participants for whom both EPImix and standard T_1_-w scans were available; additionally, some analyses focused on EPImix T_1_-w scans were repeated in the full sample of 94 participants – including the relationship between EPImix-derived GM volume and chronological age, and the correlation between EPImix-predicted age and chronological age (SI Fig. S4).

### Test-retest reliability of EPImix T_1_-w scans

Test-rest reliability of quantitative measures derived from EPImix T_1_-w scans was assessed using 10 within-session test-retest EPImix scans. Test-retest reliability was evaluated using the intraclass correlation coefficient; specifically, we used the one-way random effects model for the consistency of single measurements, i.e., ICC(3,1), hereafter referred to as ICC (Chen et al., 2018). We quantified the test-retest reliability of global tissue volumes (GM, WM and CSF), corresponding voxel-wise tissue volumes, as well as predicted brain-age.

All statistical analyses were carried out in Python 3.7.

## Results

### Tissue volume

We first compared global brain volumes of conventional and EPImix-derived T_1_-w scans. We observed strong positive correlations between standard T_1_-w and EPImix T_1_-w scans, in both GM volume (*r*_*s*_ = 0.84, p < 0.001; Fig. 1A) and WM volume (*r*_*s*_ = 0.84, p < 0.001; Fig. 1B). Measures of CSF volume demonstrated a weaker positive correlation (*r*_*s*_ = 0.56, p < 0.001; Fig. 1C). These results were qualitatively consistent when using T_1_-w scans with reduced FoV (Fig. 1D-F).

**Figure 1:**
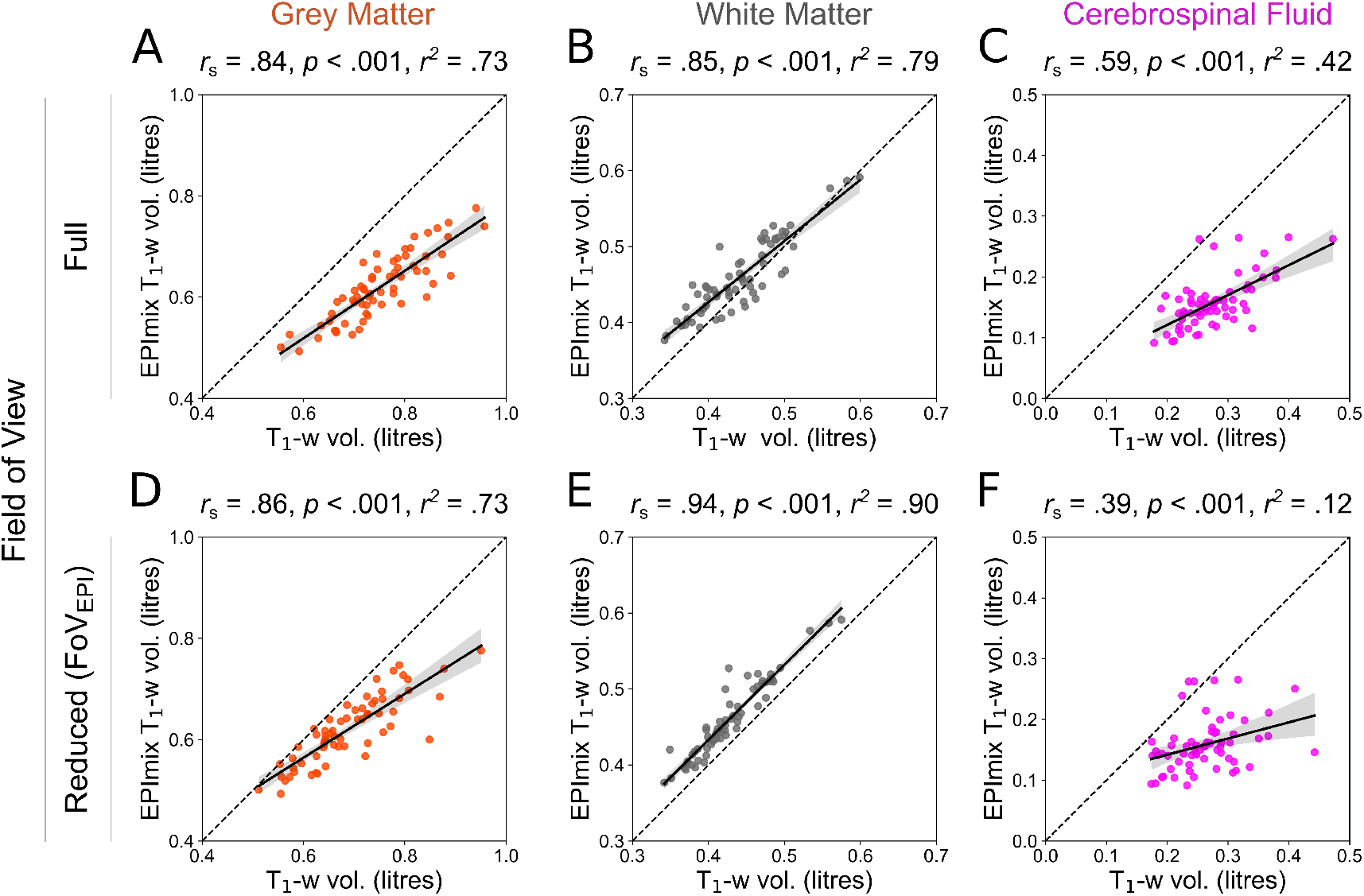
Global tissue volume estimation across contrasts. Comparison of tissue volumes (grey matter, white matter and cerebrospinal fluid) between T_1_-w and EPImix (T_1_-w) scans with full field of view (A-C) and between EPImix (T_1_-w) scans and T_1_-w scans with reduced field of view (D-F). (Spearman’s correlation coefficient *r*_*s*_, p-value, *r*^2^ derived from Pearson’s correlation.)

We next compared voxelwise estimates of GM, WM and CSF volume between conventional and EPImix-derived T_1_-w scans, in voxels with at least 0.001 mm^3^ tissue volume in at least 95% of participants. Predominantly positive correlations across participants were observed for all three tissue types. Similarly to global tissue volumes, correlations were strongest in the GM (median [1st, 3rd Quartile] (Md [Q_1_,Q_3_]) = 0.70 [0.52, 0.82]; Fig. 2A-B) and WM (Md [Q_1_,Q_3_] = 0.77 [0.62, 0.88]; Fig. 2C-D), followed by CSF (Md [Q_1_,Q_3_] = 0.53 [0.36, 0.70]; Fig. 2E-F).

**Figure 2:**
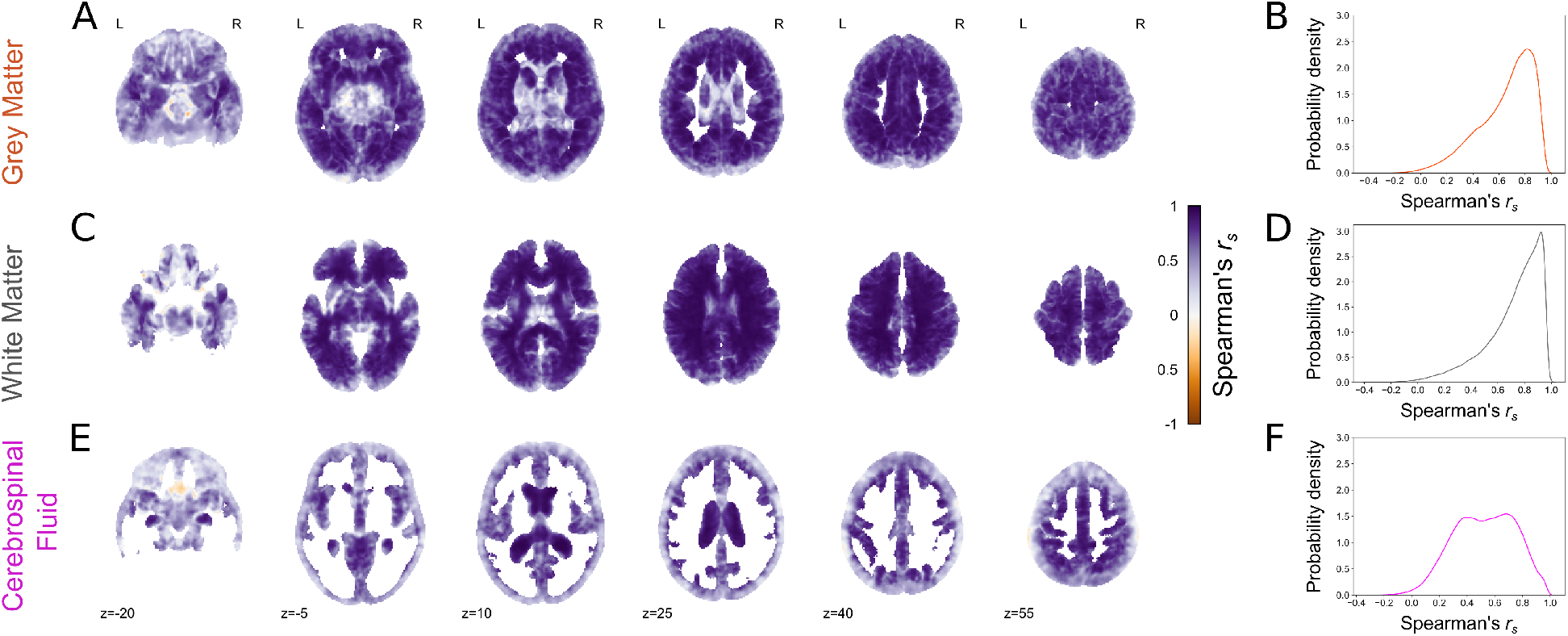
Voxel-wise tissue volume estimation across contrasts. Correlation between voxel-wise tissue volume estimates from T_1_-w and EPImix (T_1_-w) scans, and corresponding probability density plots, for grey matter (A-B), white matter (C-D) and CSF (E-F). Only voxels with at least 0.001 mm^3^ tissue volume in at least 95% of participants are shown.

Additionally, we quantified the relationship of global GM volume with age. GM volume decreased as a function of age, for estimates derived from standard T_1_-w scans (*r*_*s*_ = -0.27, p = 0.029; Fig. 3A), EPImix T_1_-w scans (*r*_*s*_ = -0.39, p < 0.001; Fig. 3B) and T_1_-w scans with reduced FoV (*r*_*s*_ = -0.27, p = 0.030; Fig. 3C).

**Figure 3:**
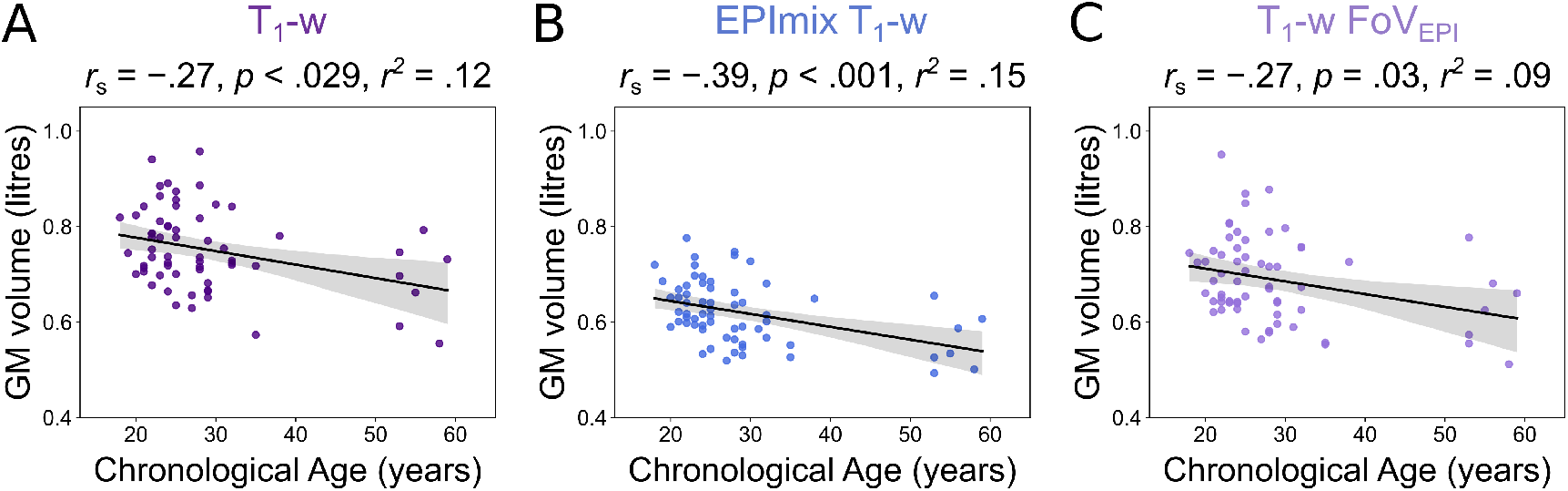
Grey matter volume as a function of chronological age, using volume estimates derived from A) T_1_-w scans, B) EPImix T_1_-w scans and C) T_1_-w scans with reduced field of view.

### Brain age

Following tissue volume analyses, we used a pre-trained Gaussian Processes Regression model (Cole, 2019) to quantify the relative ability of EPImix T_1_-w and standard T_1_-w scans to predict brain-age. As a first step, we compared the chronological (true) age of participants with their brainage prediction from each type of scan. Standard T_1_-w scans showed a high correspondence between chronological and predicted age, including both a high correlation and low error (*r*_*s*_ = 0.73, p < 0.001, MAE = 3.72 years; Fig. 4A). EPImix T_1_-w scans also showed a high correlation between chronological and predicted age (*r*_*s*_ = 0.61, p < 0.001) but with a substantially higher error (MAE = 14.20 years), due to a systematic prediction offset (Fig. 4B). Additional analysis of standard T_1_-w scans with reduced FoV (to match EPImix) showed a similarly reduced correspondence between predicted and chronological age (*r*_*s*_ = 0.36, p = 0.004, MAE = 13.05 years; Fig. 4C), suggesting that the poorer predictive ability of EPImix T_1_-w scans was likely caused by their reduced brain coverage.

**Figure 4:**
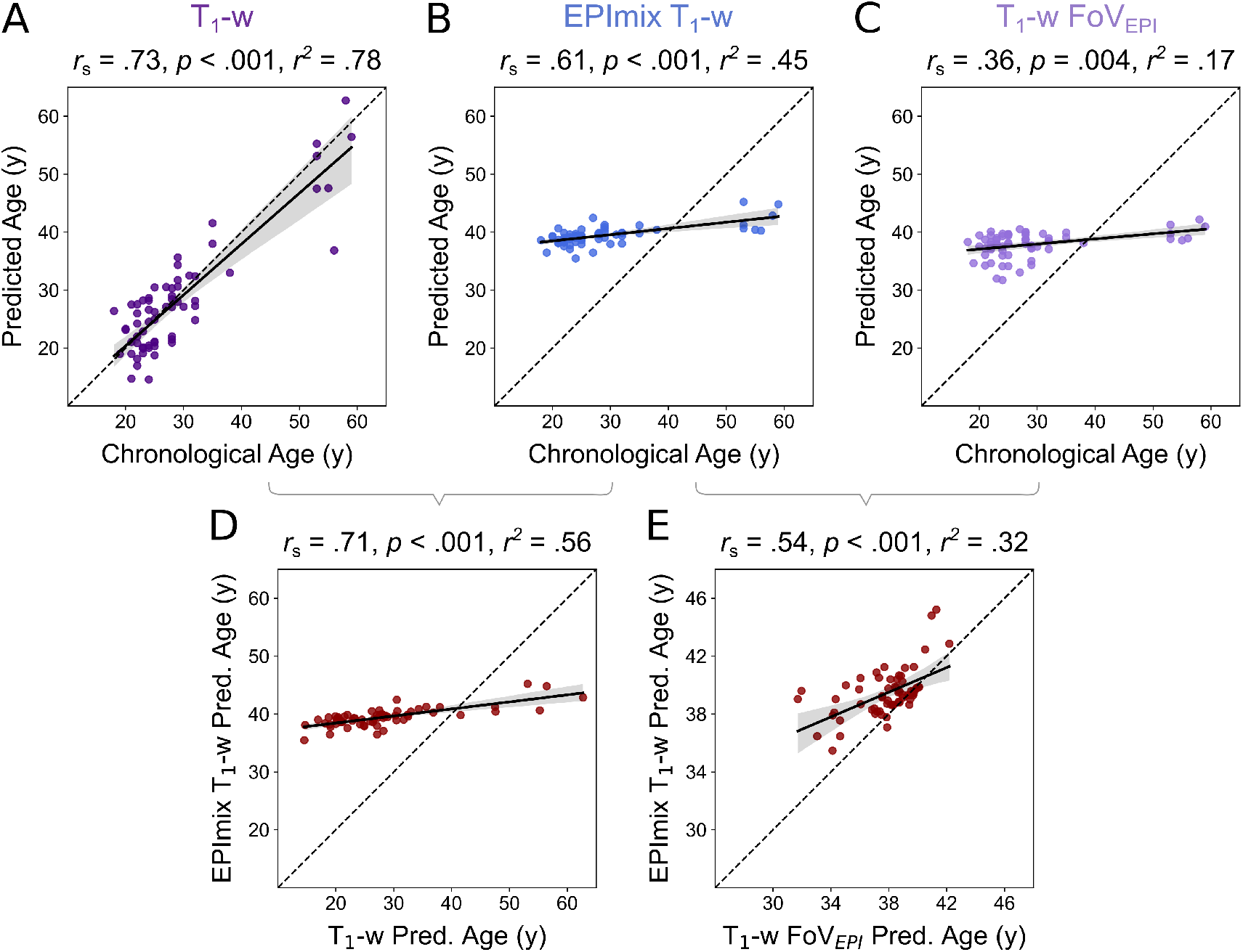
Age prediction across contrasts. Predicted age as a function of chronological age of A) T_1_-w scans, B) EPImix T_1_-w scans and C) T_1_-w scans with reduced FoV. D) Predicted age of EPImix T_1_-w scans compared to standard T_1_-w scans, and E) EPImix T_1_-w scans compared to T_1_-w scans with reduced FoV. (Spearman’s correlation coefficient *r*_*s*_, p-value, *r*^2^ derived from Pearson’s correlation.)

Furthermore, we directly evaluated the correspondence between brain-age estimates from different sequences. There was a strong positive correlation between predicted brainage derived from EPImix T_1_-w and standard T_1_-w scans (*r*_*s*_ = 0.71, p < 0.001), although the same systematic offset (relative to the identity line) described above was apparent (Fig. 4D). The correlation decreased when comparing brainage estimates from T_1_-w scans with identically reduced FoV (*r*_*s*_ = 0.54, p < 0.001), although data shifted closer to the identity line (Fig. 4E).

We next benchmarked the brain-age MAE of each contrast type relative to the worst possible MAE. Assuming an identical brain-age prediction for each participant, equivalent to the mean age of the *brainageR* training dataset (40.6 years), gives rise to an MAE of 15.6 years. While brain-age prediction using single-contrast T_1_-w scans is 4.19*×* better than this null benchmark, the performance of both EPImix T_1_-w scans and T_1_-w scans with reduced FoV is minimally improved (respectively 1.10*×* and 1.20*×*).

To improve brain-age prediction from EPImix T_1_-w scans, we used leave-one-out regression to adjust EPImix T_1_-w brain-age estimates. We regressed T_1_-w brain-age on EPImix brain-age (in 63 participants), leading to an adjusted estimate of predicted brain-age in the remaining (left-out) participant. Following iteration across all participants, this led to an adjusted MAE of 3.70 years (across left-out participants).

For all summary statistics related to predicted brainage, see Table 1.

**Table 1:** Correlation and Median Absolute Error between chronological age and predicted brain-age. The Null MAE ratio compares each MAE estimate to the MAE from the worst possible brain-age model (null MAE = 15.6 years, assuming identical prediction of the mean age of the *brainageR* training sample for all participants). The leave-one-out (LOO) cross-validation MAE was derived using regression of T_1_-w-predicted brain-age onto EPImix-predicted brain-age. For details, see *Methods* section *Statistical analysis*.

|  | EPImix T<sub>1</sub>-w | T<sub>1</sub>-w | T<sub>1</sub>-w FoV<sub>EPI</sub> |
| --- | --- | --- | --- |
| Spearman $r_s$ | 0.61 | 0.73 | 0.36 |
| Spearman p | < 0.001 | < 0.001 | 0.004 |
| $r^2$ | 0.45 | 0.78 | 0.17 |
| MAE (y) | 14.20 | 3.72 | 13.05 |
| Null MAE ratio | 1.10 | 4.19 | 1.20 |
| LOO MAE (y) | 3.70 | — | — |

### Test-retest reliability

We used a subsample of 10 participants with two (within-session) EPImix acquisitions each to quantify the test-retest reliability of all EPImix-derived quantitative measures evaluated in this study, using the intraclass correlation coefficient (ICC).

Estimates of global tissue volume showed high reliability, for GM (ICC [95% confidence interval (CI_95_)] = 0.99 [0.99,1]), WM (ICC [CI_95_] = 0.99 [0.98, 1] and CSF (ICC [CI_95_] = 0.92 [0.70,0.98]) (all p < 0.001).

Voxelwise estimates of tissue volume showed equally high test-retest reliability, for GM (Md [Q_1_,Q_3_] = 0.95 [0.90,0.97]; Fig. 5A-B), WM (Md [Q_1_,Q_3_] = 0.96 [0.90,0.98]; Fig. 5C-D) and CSF (Md [Q_1_,Q_3_] = 0.92 [0.87,0.97]; Fig. 5E-F).

**Figure 5:**
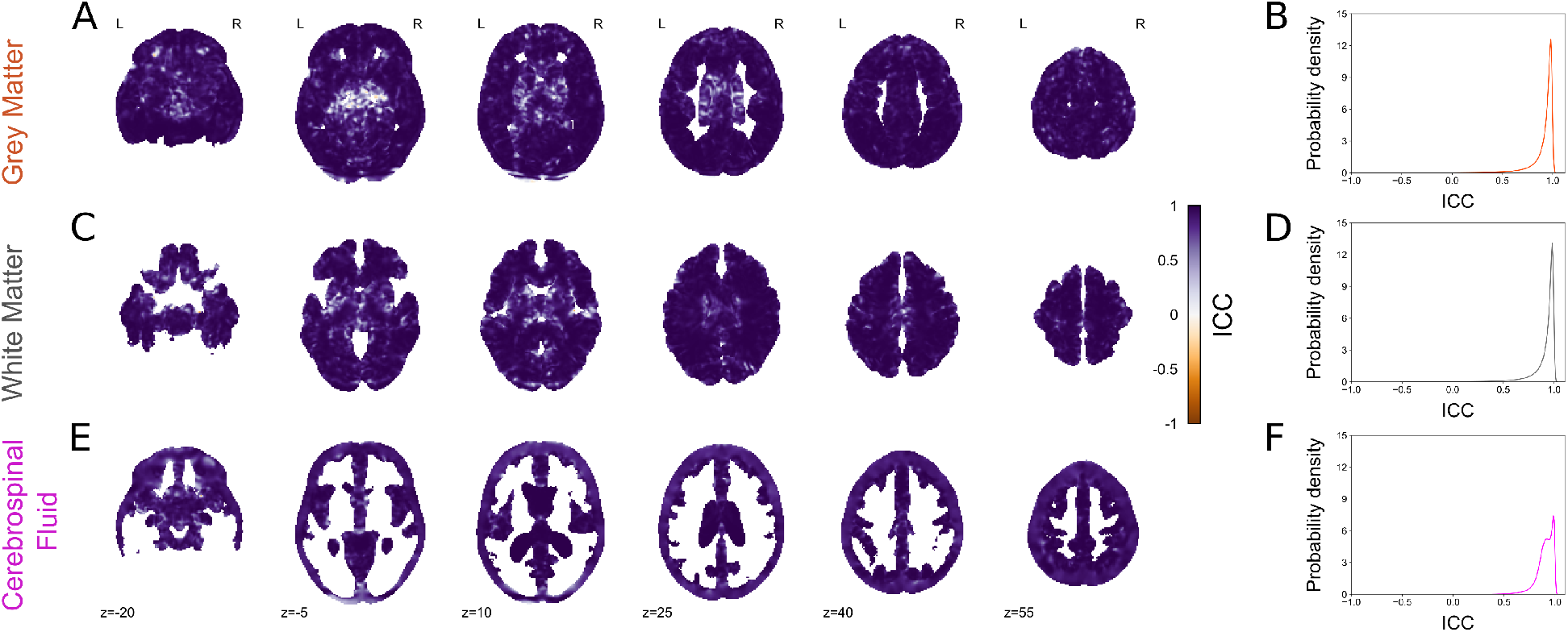
Test-retest reliability of EPImix (T_1_-w) tissue volume estimates. Reliability of voxel-wise volume estimates, and corresponding probability density plots, in grey matter (A-B), white matter (C-D) and CSF (E-F). Test-retest reliability was evaluated using the one-way random effects model for the consistency of single measurements (ICC(3,1); Chen et al. (2018)). Only voxels with at least 0.001 mm^3^ tissue volume in at least 95% of participants are shown.

Finally, brain-age estimates were equally reliable (ICC [CI_95_] = 0.99 [0.95,1]).

## Discussion

We investigated whether T_1_-w scans from the recently-developed rapid multicontrast EPImix sequence (Skare et al., 2018) are quantitatively comparable with routinely collected single-contrast T_1_-w scans. For these comparisons, we relied on interpretable and widely used quantitative measures, including global and local estimates of tissue volume as well as predicted brain-age. We found a strong correspondence between tissue volumes derived from both sequences, at both the global and local levels. Moreover, estimates derived from EPImix scans showed the expected decrease in grey matter volume as a function of age. Additionally, both types of scan significantly predicted participant brain-age, although the reduced FoV of EPImix scans led to a systematic offset and a commensurate increase in prediction error. However, we demonstrated that this can potentially be corrected using additional leave-one-out regression of T_1_-w-predicted brain-age onto EPImix-predicted brain-age. Finally, we used a subset of participants to show high test-retest reliability of all EPimix-derived quantitative measures evaluated in this study.

### Tissue volume

Both global and voxelwise analyses showed that grey matter, white matter and cerebrospinal fluid volumes are comparable between T_1_-w and EPImix T_1_-w scans. This extends our previous findings of correspondence between tissue intensities and Jacobian determinants (derived from registration of T_1_-w scans to MNI standard space), between EPImix and standard T_1_-w scans (Váša et al., 2021). While Jacobian determinants can be derived relatively rapidly from T_1_-w scans, the primary constraint being the speed of non-linear registration, they are both less interpretable and less widely used than estimates of tissue volume. For example, recent work employing the increasingly popular approach of normative modeling (Marquand et al., 2016, 2019) leveraged a large sample of scans to construct normative charts of variation in brain tissue volume across the lifespan (Bethlehem et al., 2021). Our results suggest that even volume estimates derived from rapid sequences such as EPImix could be used to anchor individual measures of brain morphology relative to such reference datasets. However, it would be preferable to increase the FoV of EPImix scans to cover the whole brain, at the cost of a small increase in acquisition time (e.g., 1.5 minutes instead of 1 minute).

In addition, healthy ageing is well known to be associated with grey matter volume reduction (Hafkemeijer et al., 2014; Ramanoël et al., 2018). Despite our modest sample size and non-uniform distribution of participants as a function of age, we found this signature of decreasing grey matter volume in EPImix T_1_-w scans.

### Predicted brain-age

Previous research has established predicted brain-age as a putative biomarker of brain health (Cole and Franke, 2017; Cole et al., 2019b). An increased predicted brainage can be a strong predictor of risk of neurodegenerative diseases, such as Alzheimer’s disease, and neuropsychiatric disorders, such as schizophrenia (Cole and Franke, 2017; Franke and Gaser, 2019; Cole, 2020). We used a pre-trained Gaussian Processes Regression model (Cole, 2019) to derive estimates of predicted brain-age from EPImix as well as conventional T_1_-w scans. EPImix-derived brain-age showed a strong correspondence (as quantified using Spearman’s *r*_*s*_, and *r*^2^) to chronological age – of a similar magnitude to previous studies (e.g. Cole, 2020) – as well as to brain-age estimated from standard T_1_-w scans.

However, a systematic offset led to a large median absolute error of the prediction. Re-analysis of T_1_-w scans with reduced FoV led to a commensurate drop in performance, confirming that this systematic error is likely caused by incomplete brain coverage of EPimix scans, combined with the fact that the *brainageR* model we used (Cole, 2019) was pre-trained on conventional single-contrast T_1_-w scans (with full FoV). However, we showed that this systematic offset can potentially be corrected using leave-one-out regression of T_1_-w-predicted brain-age on EPImix-predicted brain-age, to yield an adjusted estimate of EPImix-derived brain-age. Such approaches could be used to adjust brain age predictions derived from newly-acquired EPImix scans with reduced FoV.

In future, multicontrast data produced by the EPImix sequence could also be used to improve the accuracy of brain age predictions, as demonstrated by prior work (Cole, 2020).

### Active acquisition

An exciting potential application for the EPImix sequence is the active acquisition approach proposed by Cole et al. (2019a). Active acquisition involves the online analysis of MR scans, aiming to use active learning (Settles, 2009) to analyse scans as they are being acquired, in turn guiding subsequent acquisition steps (Cole et al., 2019a). As our study has found comparable quantitative measures between EPImix and routinely collected T_1_-w scans, whilst scanning times are considerably faster, there is potential for EPImix to be utilised in the online collection and analysis of brain scans, in the process of active acquisition.

Cole et al. (2019a) proposed three examples of active acquisition scenarios, with EPImix potentially being suitable for use in all three. Firstly, due to its relative speed, EPImix could be used to rapidly acquire low resolution data to inform whether higher resolution data needs to be acquired (more slowly), depending on detection of abnormalities in the EPImix scan. The other two scenarios propose to leverage multi-modal data, to classify participants and/or to identify modalities in which participants deviate the most from the norm. Due to the multicontrast nature of the EPImix sequence, it could also be used in such scenarios. However, a larger normative sample would be required to robustly model inter-individual variability, and clinical data would be useful to test the translational relevance of such models (Marquand et al., 2016, 2019).

The acquisition time of the EPImix sequence is considerably faster than analogous single-contrast scans (Skare et al., 2018); however, the current analysis pipeline (median = 4 min 49 seconds) is too slow for real-time analysis required for active acquisition (Cole et al., 2019a). As it currently stands, a small amount of analyses could be carried out whilst the participant is in the scanner, to inform several additional acquisition steps. Although not sufficient for the near-real-time process of active acquisition, this could still reduce the need for participants to be recalled for follow-up scans.

In the future, there is potential to run the data analysis on more powerful computers, to speed up the processing and analysis pipeline. Alternatively, deep learning tools hold the promise of massively accelerating processing. While such models are very costly to train – in terms of time, computing power and energy – inference is very fast. Examples of relevant recent tools include *SynthSeg* for scan segmentation (Billot et al., 2020), *voxelmorph* for scan registration (Hoffmann et al., 2020) or *FastSurfer*, a deep learning analogue of *Freesurfer* which is particularly suitable for surface-based measures (Henschel et al., 2020). Additionally, EPImix scans could be super-resolved using image quality transfer tools, to improve scan resolution and likely increase correspondence to conventional single-contrast acquisitions (Iglesias et al., 2021).

### Methodological considerations

There are a number of methodological limitations to be considered, to maximise potential practical utility of the rapid EPImix sequence in the context of quantitative analysis.

One limitation of the present study is the reduced FoV of EPImix scans, particularly in outer areas of cortex, such as the inferior temporal lobe and superior parietal lobe. We investigated the impact of this reduced coverage on our results by re-analysing single-contrast T_1_-w scans with identically reduced FoV (on an individual participant basis). This enabled us to conclude that the reduced FoV is likely the primary hindrance in the ability of EPImix T_1_-w scans to predict brain-age. However, this issue can potentially be addressed using an additional leave-one-out regression step to correct predicted brain-age values, as demonstrated here. We relied on leave-one-out cross-validation – despite its inflated performance relative to k-fold or hold-out validation – due to small sample size, and to demonstrate the utility of even a small paired (EPImix and single-contrast T_1_-w scan) dataset for the correction of brain-age predictions in a new (EPImix-only) scan. An improved approach in future studies would be to increase the FoV of EPImix scans; despite increased scanning time, the multicontrast EPImix sequence would still remain considerably faster than routinely collected (higher-resolution) single-contrast scans. A larger FoV could be particularly important in clinical settings, where accurate diagnosis is dependent on comprehensive brain coverage (Ou et al., 2018).

In addition, the image analysis pipeline used here remains too slow to realistically be used in an active acquisition setting, particularly if image processing needs to occur in near-real-time to drive scanning in a closed loop (Cole et al., 2019a). Our previous work developed a custom minimal processing pipeline which is sufficiently fast to realistically operate in near-real-time; however, this is limited to scan registration and Jacobian extraction, and does not enable quantification of tissue volume or brain age prediction. Increasing the speed of processing would be particularly valuable to enable the EPImix sequence to be used as a quantitative screening test, given that its speed is one of its main advantages over standard structural MR imaging (Skare et al., 2018; Delgado et al., 2019). One potential avenue for the creation of custom rapid processing pipelines is Bayesian optimisation, which has previously been used to customise processing tools for brain age prediction (Lancaster et al., 2018).

Moreover, additional research is needed to quantitatively compare the other contrasts generated by the multicontrast EPImix sequence, including T_2_-weighted, T_2_-FLAIR, T_2_*-weighted, diffusion-weighted contrasts and the apparent diffusion coefficient, with analogous single-contrast MR scans. Similarly, it would be interesting to compare EPImix to other similar sequences on scanners from other manufacturers (Polak et al., 2020), using quantitative tools applied here as well as the aforementioned deep-learning approaches.

Finally, further research should focus on clinical groups. Beyond the translational value of such studies, they would help to further investigate the correspondence between EPImix and standard T_1_-w scans. In particular, a key question is whether group differences are similar within EPImix and standard T_1_-w scans.

### Conclusion

In summary, we used popular quantitative measures derived from brain MRI to compare T_1_-w scans derived from the new rapid EPImix sequence with routinely collected single-contrast T_1_-w scans. We found that both global and voxelwise tissue volume estimates derived from EPImix T_1_-w scans were comparable with analogous measures extracted from single-contrast high-resolution T_1_-w scans.

Brain age predictions from EPImix T_1_-w scans showed a strong correlation with both chronological age and brainage estimates from single-contrast T_1_-w scans, although a systematic offset led to a high prediction error. However, this could be corrected using additional leave-one-out regression.

Taken together, our findings underline the potential of the EPImix sequence to reduce scanning time, increasing participant comfort and reducing cost, and further highlight its relevance and applicability to quantitative MR analysis routines. Future extension of this work includes the development of additional customised and deep-learning-based processing tools, as well as the analysis of other contrasts generated by EPImix; this will enable researchers to fully harness the potential of this multicontrast sequence to drive innovative paradigms in MRI acquisition and processing.

## Availability of code and data

All processing and analysis code is available on FV’s GitHub, at https://github.com/frantisekvasa/epimix_volume_brain_age. Processed EPImix and single-contrast T_1_-w data, including voxel-wise tissue volumes and brain age estimates, are available at https://doi.org/10.6084/m9.figshare.18128225 (Váša, 2022).

## Acknowledgments

FV & RL would like to acknowledge support from the Data to Early Diagnosis and Precision Medicine Industrial Strategy Challenge Fund, UK Research and Innovation (UKRI). RL received support from the Wellcome/EPSRC Centre for Medical Engineering (WT 203148/Z/16/Z). We would also like to acknowledge support from NIHR Maudsley Biomedical Research Centre (BRC) NNA08. JC acknowledges funding from a UKRI/MRC Innovation Fellowship (MR/R024790/2).

## Footnotes

1 In this context, the terms *quantitative* and *qualitative* refer to the type of analysis used, rather than the type of MR scan. Specifically, qualitative measures refer to visual inspection of scans, whereas quantitative measures refer to analyses which employ numerical computations.

2 Note that the EPImix sequence includes a T_1_-FLAIR contrast, while the “conventional” single-contrast scans were acquired using an IR-FSPGR sequence. Still, as both sequences are T_1_-weighted, we refer to both as such (as well as simply “T_1_-w”).

