## Supplementary Information for "Tissue volume estimation and age prediction using rapid structural brain scans"

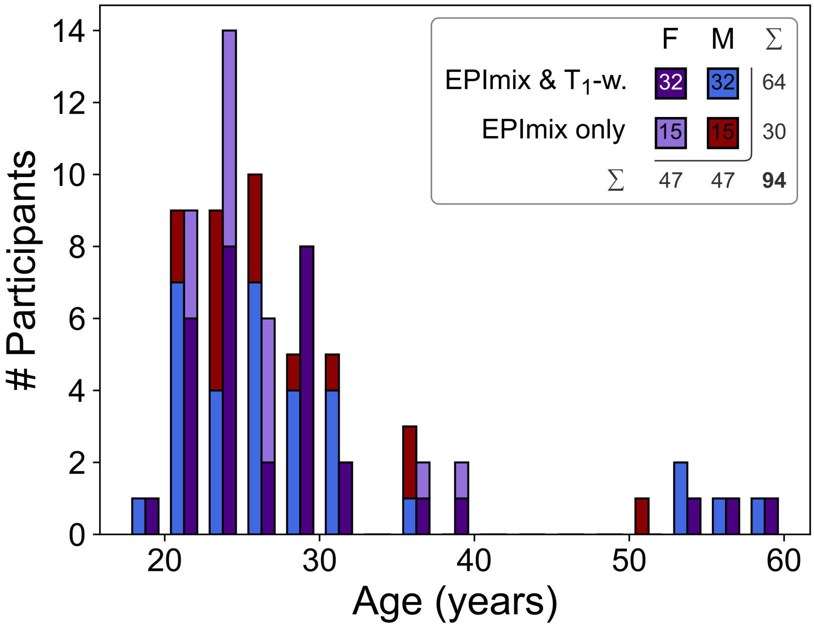


**Figure S1: Age distribution of participants by sex and scan sequence.** Scans from a total of 94 participants (47 female, 47 male) were included in this study. Of those, 64 (32 female, 32 male) were scanned using both EPImix and single-contrast $T_1$-w sequences, while an additional 30 (15 female, 15 male) were scanned using EPImix only. There were no differences in participant numbers by sex or scan sequence (Chi-squared test, $\chi^2 = 0$, P = 1).

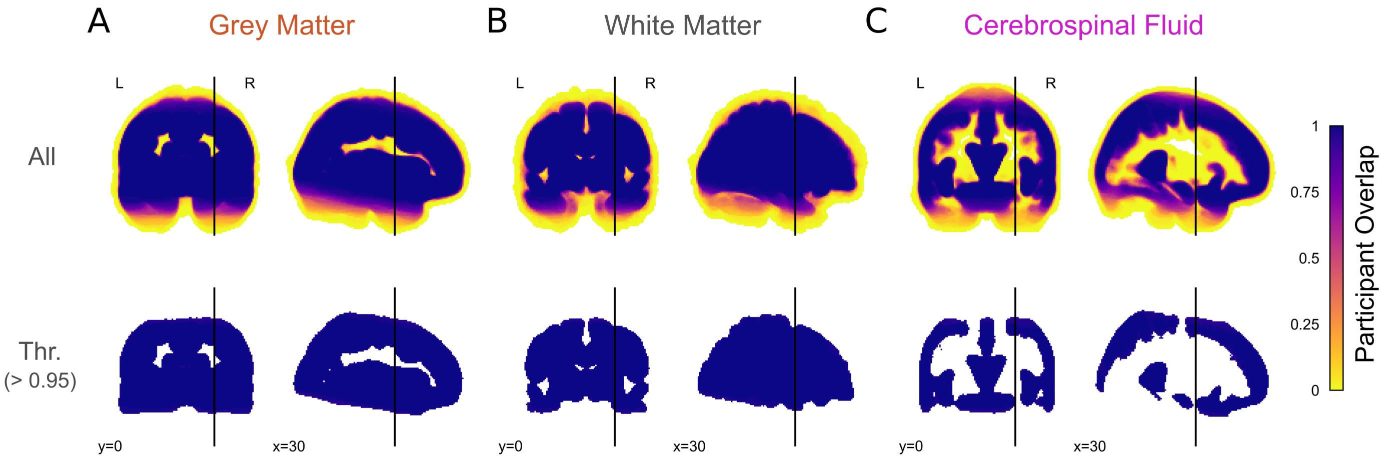


**Figure S2: Participant overlap for each tissue type in EPImix scans with reduced FoV.** Top row: Proportion of participants with at least 0.001 $mm^3$ tissue volume at each voxel of A) grey matter, B) white matter and C) cerebrospinal fluid. Subsequent analyses were limited to voxels with at least 95% participant overlap (bottom row).

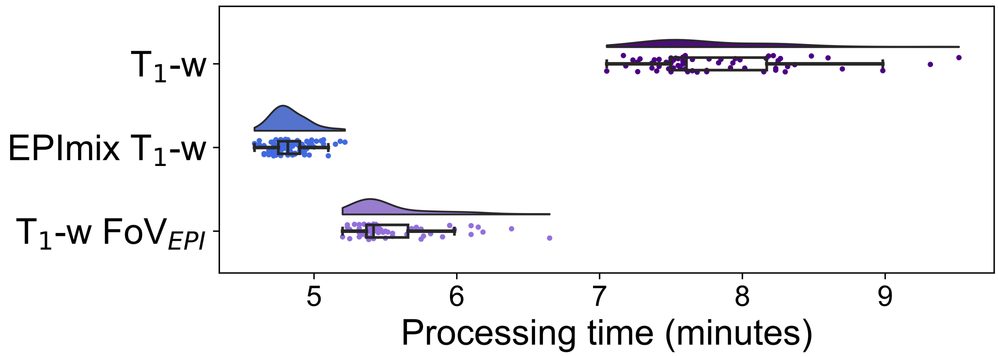


**Figure S3: Total processing time (in minutes)** for standard $T_1$-w scans, EPImix $T_1$-w scans and $T_1$-w scans with reduced field-of-view.

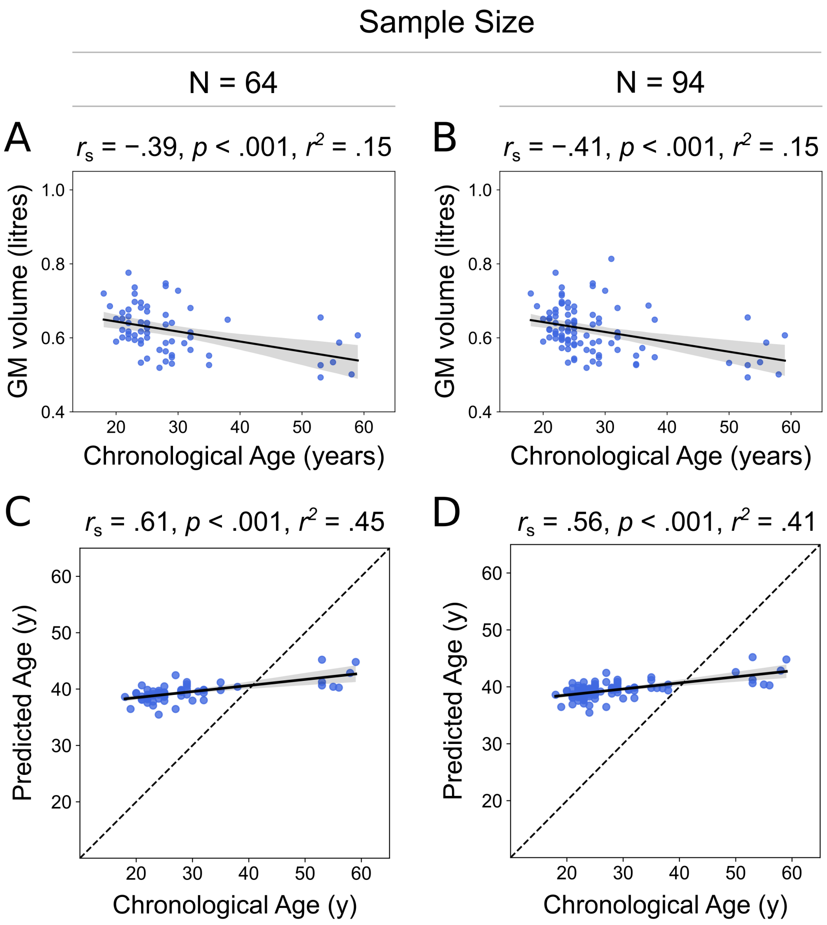


**Figure S4: Consistency of EPImix results across sample sizes.** Top row: GM volume as a function of chronological age, in A) the subset of 64 participants with both EPImix and single-contrast $T_1$-w scans, and B) the full sample of 94 participants with EPImix scans. Bottom row: Predicted age as a function of chronological age, in C) the subset of 64 participants with both EPImix and single-contrast $T_1$-w scans, and D) the full sample of 94 participants with EPImix scans. (Note that panel A is identical to main text Figure 3B, and panel C is identical to main text figure 4B.)
